# Pharmacogenetics of antidepressant response: a polygenic approach

**DOI:** 10.1101/093799

**Authors:** Judit García-González, Katherine E. Tansey, Joanna Hauser, Neven Henigsberg, Wolfgang Maier, Ole Mors, Anna Placentino, Marcella Rietschel, Daniel Souery, Tina Žagar, Piotr M. Czerski, Borut Jerman, Henriette N. Buttenschøn, Thomas G. Schulze, Astrid Zobel, Anne Farmer, Katherine J. Aitchison, Ian Craig, Peter McGuffin, Michel Giupponi, Nader Perroud, Guido Bondolfi, David Evans, Michael O’Donovan, Tim J. Peters, Jens R. Wendland, Glyn Lewis, Shitij Kapur, Roy Perlis, Volker Arolt, Katharina Domschke, Major Depressive Disorder Working Group of the Psychiatric Genomic Consortium, Gerome Breen, Charles Curtis, Lee Sang-Hyuk, Carol Kan, Stephen Newhouse, Hamel Patel, Bernhard T. Baune, Rudolf Uher, Cathryn M. Lewis, Chiara Fabbri

**Author notes:** These two authors jointly supervised the study. **Corresponding author:** Professor Cathryn Lewis, PhD, Social, Genetic and Developmental Psychiatry Centre, Institute of Psychiatry, Psychology & Neuroscience, De Crespigny Park, London SE5 8AF.

## Abstract

**Background:** Major depressive disorder (MDD) has a high personal and socio-economic burden and more than 60% of patients fail to achieve remission with the first antidepressant. The biological mechanisms behind antidepressant response are only partially known but genetic factors play a relevant role. A combined predictor across genetic variants may be useful to investigate this complex trait.

**Methods:** Polygenic risk scores (PRS) were used to estimate multi-allelic contribution to: 1) antidepressant efficacy; 2) its overlap with MDD and schizophrenia. We constructed PRS and tested whether these predicted symptom improvement or remission from the GENDEP study (n=736) to the STAR*D study (n=1409) and vice-versa, including the whole sample or only patients treated with escitalopram or citalopram. Using summary statistics from Psychiatric Genomics Consortium for MDD and schizophrenia, we tested whether PRS from these disorders predicted symptom improvement in GENDEP, STAR*D, and five further studies (n=3756).

**Results:** No significant prediction of antidepressant efficacy was obtained from PRS in GENDEP/STAR*D but this analysis might have been underpowered. There was no evidence of overlap in the genetics of antidepressant response with either MDD or schizophrenia, either in individual studies or a meta-analysis. Stratifying by antidepressant did not alter the results.

**Discussion:** We identified no significant predictive effect using PRS between pharmacogenetic studies. The genetic liability to MDD or schizophrenia did not predict response to antidepressants, suggesting differences between the genetic component of depression and treatment response. Larger or more homogeneous studies will be necessary to obtain a polygenic predictor of antidepressant response.

## Introduction

Major depressive disorder (MDD) is a common mental disorder characterized by sadness, anhedonia, guilt, feelings of low self-worth, poor concentration, disturbed appetite and sleep and suicidal thoughts (World Health Organization, 1993; American Psychiatric Association, 2013). Its heavy socio-economic and individual burden makes it a global concern: lifetime prevalence of MDD ranges from 10% to 15% and MDD is one of the top ten causes of years lived with disability (YLDs) worldwide (The WHO World Mental Health Survey Consortium, 2004; Global Burden of Disease Study 2013 Collaborators, 2015).

Antidepressant drugs are the first-line treatment for MDD, with more than 30 antidepressant drugs available (Fabbri *et al.*, 2016). Responses vary widely across individuals: one third of patients show complete remission after the first drug prescribed, one third improves after a change of treatment or augmentation, and one third fail to respond after two different antidepressants prescribed (Trivedi *et al.*, 2006; Souery *et al.*, 2011). For each patient, the most effective treatment can only be identified by trial and error - a lengthy process which delays recovery and leads to poorer clinical outcomes (Steimer *et al.*, 2001). The ability to identify the most effective drugs for each patient or to predict treatment resistance would be a turning point in MDD treatment, enabling personalized prescribing. However, no predictor of antidepressant response is currently available; clinical characteristics are weak predictors of improvement in depressive symptoms, and no established biomarkers or genetic signatures exist for antidepressant response.

Genome-wide association studies (GWAS) to identify single nucleotide polymorphisms (SNPs) associated with antidepressant response have provided tentative hints, but most associations have been inconclusive and are unreplicated (Myung *et al.*, 2015; Sasayama *et al.*, 2013; Biernacka *et al.*, 2015; GENDEP Investigators, MARS Investigators and STAR*D Investigators, 2013; Uher *et al.*, 2010; Trivedi *et al.*, 2006; Ising *et al.*, 2009). These disappointing findings may be ascribed to several features of pharmacogenetic studies: limited sample size, heterogeneity between studies in design, drug, and assessment of outcome. Given the challenges of accruing sufficiently strong evidence to confirm association of a single SNP with antidepressant response, an alternative approach is to construct a single summary genetic variable representing genome-wide information which can be used for prediction.

Polygenic risk scores (PRS) capture in a single variable the additive effect of SNP alleles across the genome (Dudbridge, 2013). In contrast to GWAS analysis, where a single SNP must reach stringent significance levels, PRS are constructed from multiple SNPs with lower evidence of association, with the assumption that genetic markers that do not meet the genome-wide significance threshold might have good predictive power when they are considered collectively.

In this study we test whether polygenic risk scores can provide prediction of response to anti-depressants, building PRS directly from clinical trials of antidepressant response (STAR*D, GENDEP) (Garriock *et al.*, 2010; Uher *et al.*, 2010), and secondly testing the hypothesis of whether genetic liability to the psychiatric disorders of MDD and schizophrenia contributes to variation in antidepressant response. Indeed an overlap between the genetics of MDD and antidepressant response has been hypothesized, but MDD also shares susceptibility genetic factors with schizophrenia (Cross-Disorder Group of the Psychiatric Genomics Consortium, 2013), suggesting a possible overlap between the genetics of schizophrenia and antidepressant response. We analyse two large pharmacogenetic trials (GENDEP, STAR*D) and expand our study to other studies of antidepressant response, giving a substantial sample size in which to develop and test predictors of treatment response.

## Materials and Methods

### Pharmacogenetic studies

Seven pharmacogenetic studies were included, all similar in their fundamental features: (1) participants were treatment-seeking individuals diagnosed with MDD based on DSM-IV/ICD-10 criteria (World Health Organization, 1993; American Psychiatric Association, 2013), with other psychiatric diagnoses excluded (schizophrenia spectrum disorders, bipolar disorders, current alcohol or drug dependence). For each study participant, prospective data on outcome of antidepressant treatment were recorded according to standard and comparable scales. Missing end-point measurements were imputed using the best unbiased estimate from a mixed-effect linear regression model, with fixed linear and quadratic effects of time and random effects of individual and centre of recruitment (for multi-centric studies) according to previous studies (Tansey *et al.*, 2012; Uher *et al.*, 2010). Patients were included in the analyses only if baseline assessment and at least one post-baseline assessment were available.

The GENDEP and STAR*D studies formed our primary studies for discovery and testing variants specific to antidepressant response. For testing the hypothesis that genetic liability for MDD and schizophrenia predicts antidepressant response, we included four further trials from the NEWMEDS consortium (GENPOD, GODS, GSK, Pfizer) (Tansey *et al.*, 2012) and a newly genotyped naturalistic study from the University of Muenster (Baune *et al.*, 2008, 2010). All studies were approved by local ethics boards of participating centres, and all participants provided written informed consent after the study procedures were explained and prior to sample collection. Detailed information for each sample are given in Table 1 and Supplementary Methods.

**Table 1.**
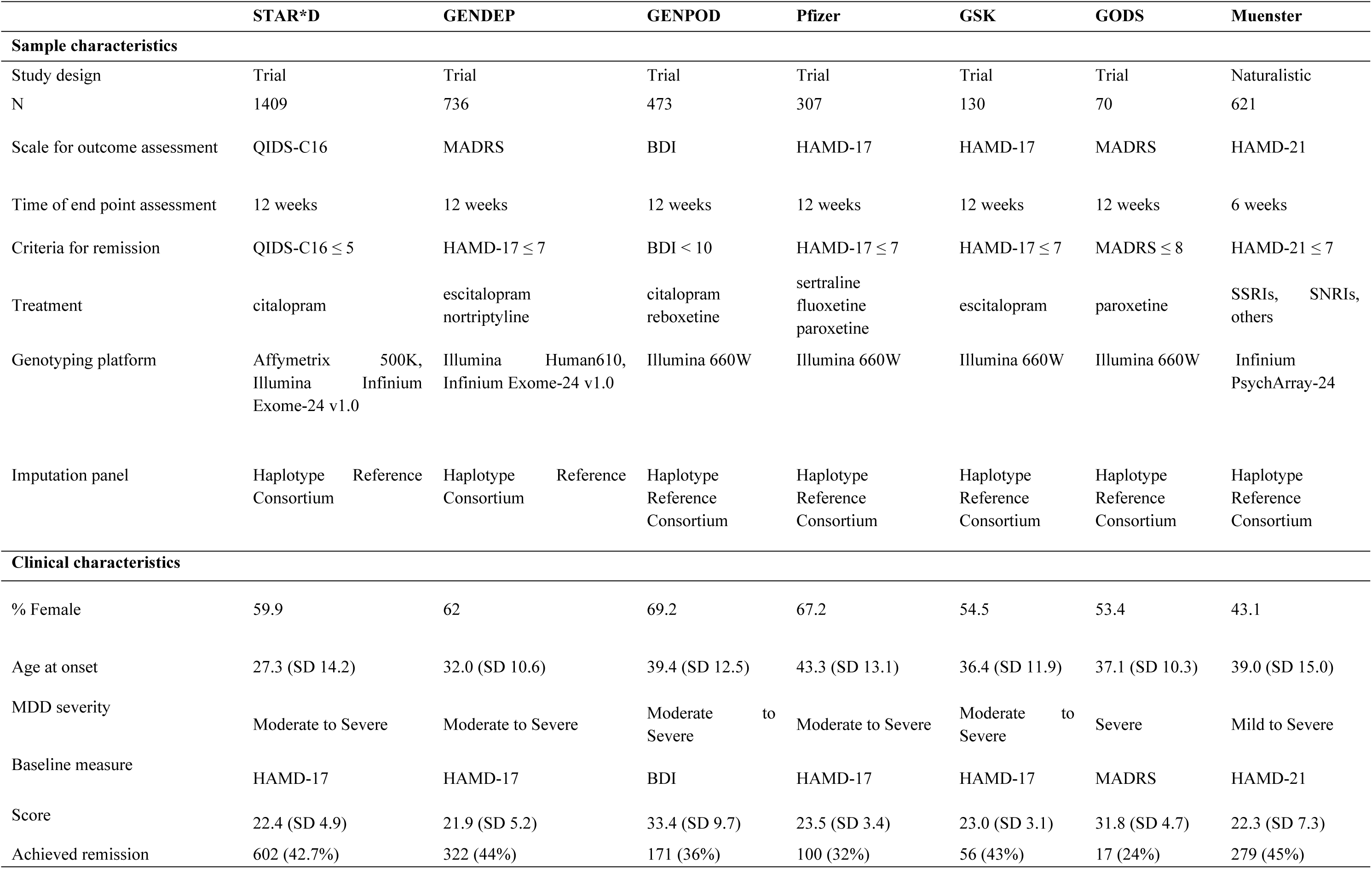
Pharmacogenetic study characteristics. MADRS; Montgomery-Åsberg Depression Rating Scale. QIDS-C16; Quick Inventory of Depressive Symptomatology-Clinician Rated. BDI; Beck Depression Inventory. HAMD-17; Hamilton Rating Scale for Depression (17 items). HAMD-21; Hamilton Rating Scale for Depression (21 items). SSRIs; selective serotonin reuptake inhibitors. SNRIs; serotonin and noradrenaline reuptake inhibitors.

### Outcome measures

Two phenotypes were investigated at the end-point of each study, a continuous measure of improvement, calculated as the percentage change in symptom score, and symptom remission (Table 1). Percentage change was preferred to absolute change because it is less correlated with initial severity, relatively independent of the scale, and closely reflects clinician’s impression of improvement (Uher *et al.*, 2009; Lane, 2008; Mallinckrodt, Clark and David, 2001). Remission was defined as a score below a consensus cut-off that corresponds to absence of depression for each scale (Hamilton, 1967; Montgomery and Asberg, 1979; Beck *et al.*, 1961). For GENDEP, remission was defined using HAMD-17, since there was stronger consensus about the threshold to identify remission on this scale compared to MADRS (Uher *et al.*, 2008). Remission has lower power to detect an effect than a continuous measure (Streiner, 2002) but it may be associated with MDD prognosis (Gaynes *et al.*, 2009).

### Psychiatric Genomics Consortium summary statistics

Genome-wide summary statistics for large meta-analysis from the Psychiatric Genomics Consortium (PGC) were used to construct PRS for MDD and schizophrenia for each participant in the pharmacogenomic studies. Summary statistics for schizophrenia were downloaded from pgc.unc.edu (36,989 schizophrenia cases, 113,075 controls) (Ripke *et al.*, 2014). MDD summary statistics were from the latest PGC MDD meta-analysis comprising 51,865 MDD cases and 112,200 controls (unpublished data).

### Statistical analysis

Individual-level genotypes were available for all pharmacogenetic studies. GENDEP and STAR*D were imputed using genotype data from genome-wide and exome arrays capturing both common and rare variation (Table 1; Supplementary Methods). The Muenster study was imputed from Infinium PsychArray-24, and phenotype and genotype data from studies in the NEWMEDS consortium (GENPOD, Pfizer,GSK, GODS) were used as previously reported (Tansey *et al.*, 2012). All these studies were imputed using Minimac3 and the Haplotype Reference Consortium (HRC version 1) as reference panel. In STAR*D and GENDEP, tests of SNP association were performed using linear regression (for percentage change in symptom score) and logistic regression (remission) using PLINK (Purcell *et al.*, 2007). Each model included covariates of ancestry-informative principal components, age, baseline severity of depression and ascertainment centre (for multi-centre studies of STAR*D, GENDEP and Pfizer).

GWAS summary data from GENDEP, STAR*D, PGC-SCZ, and PGC-MDD were used as discovery studies. A schematic representation of study design is provided in Figure 1. SNPs were clumped by linkage disequilibrium (LD) and p-value: SNP with the smallest p-value within a 250 Kb window were retained, and all SNPs in LD (r^2^ > 0.1) with the retained SNP were excluded. When PGC-MDD was used as discovery study, markers with allele frequency difference of over 0.15 between discovery and test data sets were excluded to ensure comparability given the different genotyping chips and imputation reference panels used. PRS were constructed using the software PRSice v.1.25 (Euesden, Lewis and O’Reilly, 2015). PRS were calculated as the sum of associated alleles, weighted by effect sizes, for SNPs with p-values less than pre-defined threshold P_T_. Nine p-value thresholds of P_T_ < (0.0001, 0.001, 0.05, 0.1, 0.2, 0.3, 0.4, 0.5, 1) were used with all pruned SNPs included in the final threshold P_T_=1. Symptom improvement and remission outcomes were regressed on polygenic scores, adjusting for the covariates as used in the GWAS analyses, and compared to a model including only covariates. The proportion of phenotypic variance explained by PRS was assessed by R^2^ (for improvement) or Nagelkerke’s R^2^ (for remission). To decrease pharmacological heterogeneity across samples and to increase power, analyses were repeated stratifying by antidepressant, including only studies using escitalopram and citalopram (STAR*D, GSK, 417 GENDEP participants, 242 GENPOD participants and 121 Muenster participants).

**Figure 1:**
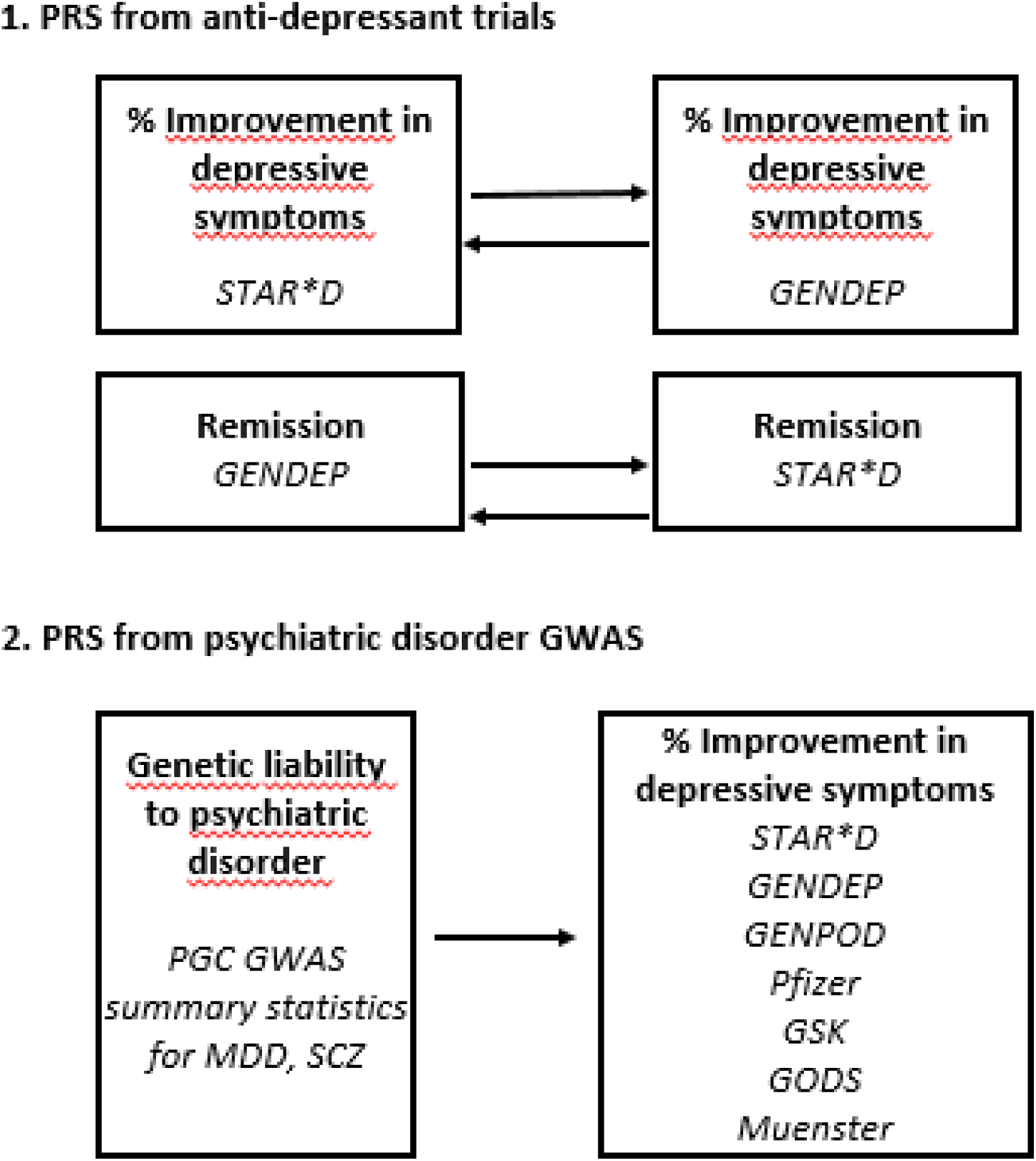
Study design, capturing (1) prediction of improvement and remission using large antidepressant response trials as discovery studies, and (2) prediction from psychiatric disorder PRS, into all antidepressant studies. Arrows indicate PRS from discovery to test data sets. PGC=Psychiatric Genomics Consortium.

Prediction of improvement from MDD and schizophrenia was implemented separately in each pharmacogenetic study, then a fixed effects meta-analysis was performed to combine results across studies at each P_T_.

A Bonferroni correction was applied to account for multiple testing. We estimated p=0.01 as an approximate correction for correlation between PRS at 9 PT values, and then corrected further for four independent hypotheses, giving a required significance level of p=0.0025.

### Power calculation

Power calculations for the polygenic analysis were performed using the R package AVENGEME (Palla and Dudbridge, 2015), at each P_T_. Models assumed SNP heritability of 0.21 for MDD (Cross-Disorder Group of the Psychiatric Genomics Consortium *et al.*, 2013), 0.42 for response to antidepressants (Tansey *et al.*, 2013) and 0.33 for schizophrenia (Ripke *et al.*, 2013a). Lifetime prevalences used were 16.2% for MDD (Kessler *et al.*, 2003), and 0.87% for schizophrenia (Perälä *et al.*, 2007). The models used for power calculation assumed that the markers are independent and 5% of SNPs have an effect in the training phenotype. For cross-trait polygenic analysis (MDD, schizophrenia and antidepressant response), two hypothetical scenarios were tested, comparing change in prediction accuracy when covariance between genetic effects in the training and target samples were 25% or 50%.

With GENDEP or STAR*D as discovery sample, the power to detect the genetic contribution of response to antidepressants was limited (12% for improvement, 8% for remission). Using PGC MDD and PGC SCZ as discovery had higher power. Assuming a covariance of 25% between SCZ and improvement in depression symptoms gave >90% power in the combined pharmacogenetic samples. A covariance of 50% between MDD and improvement in depression symptoms had 90% power to detect an effect in the combined pharmacogenetic sample, but only power of 37% with 25% covariance.

## Results

Firstly, we tested whether PRS predict improvement and remission in depression symptoms after twelve weeks of antidepressant treatment, using GENDEP and STAR*D. Each study was used as discovery and then as target study, testing the PRS constructed from STAR*D GWAS results in GENDEP, and vice-versa. No significant prediction of treatment response was attained for improvement or for remission in the whole sample (Supplementary Figure 1) or restricting the analysis to citalopram/escitalopram (Supplementary Figure 2) treated patients (Table 2). The lowest p-value of p=0.023, using the GENDEP remission GWAS to predict remission in STAR*D, did not reach the required Bonferroni correction of 0.0025.

**Table 2:**
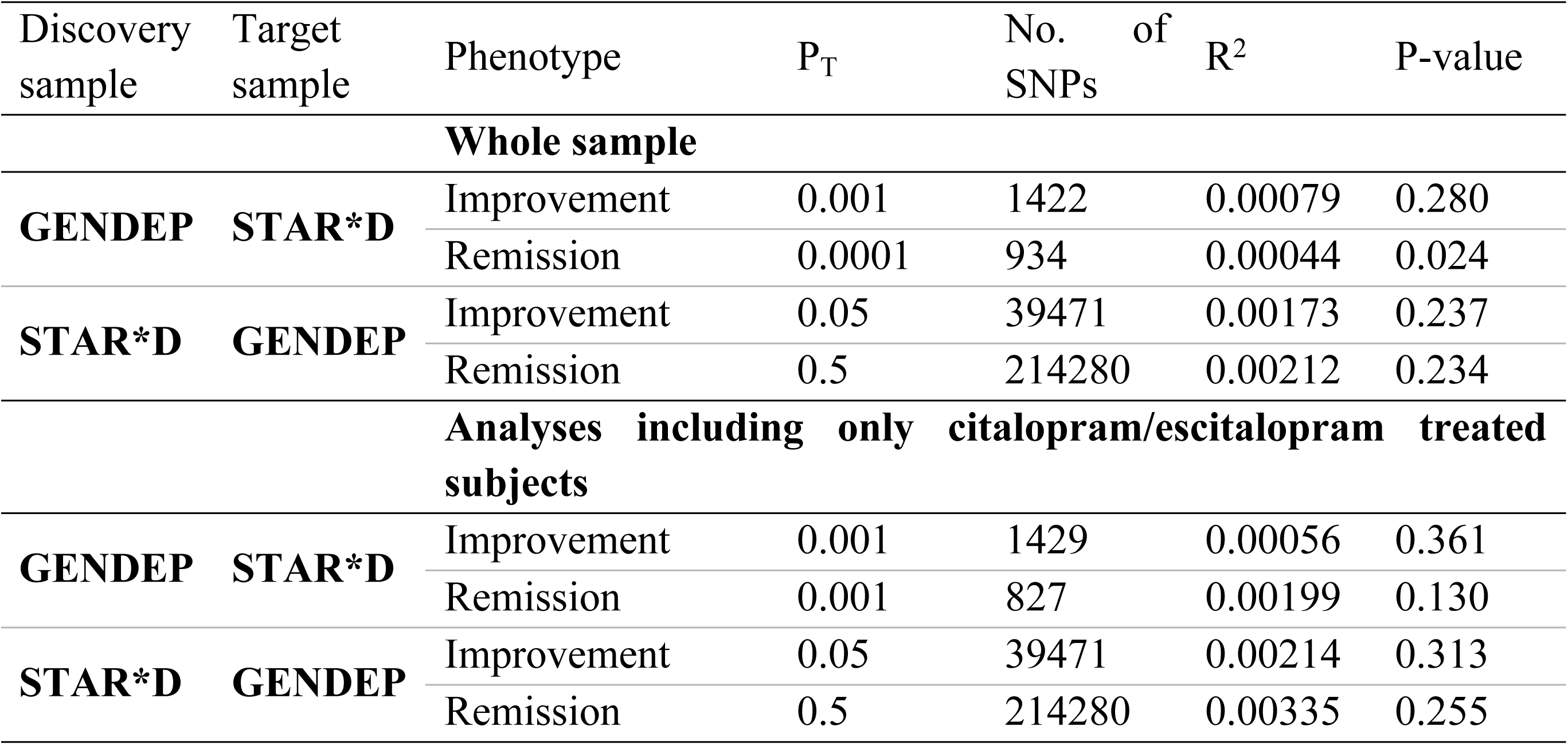
Prediction of improvement in depression symptoms and remission after 12 weeks of antidepressant treatment. Results are shown for the P_T_ threshold attaining the lowest p-value. R^2^; Proportion of variance explained.

| Discovery sample | Target sample | Phenotype | P<sub>T</sub> | No. of SNPs | R<sup>2</sup> | P-value |
| --- | --- | --- | --- | --- | --- | --- |
| Whole sample |  |  |  |  |  |  |
| GENDEP | STAR*D | Improvement | 0.001 | 1422 | 0.00079 | 0.280 |
|  |  | Remission | 0.0001 | 934 | 0.00044 | 0.024 |
| STAR*D | GENDEP | Improvement | 0.05 | 39471 | 0.00173 | 0.237 |
|  |  | Remission | 0.5 | 214280 | 0.00212 | 0.234 |
| Analyses including only citalopram/escitalopram treated subjects |  |  |  |  |  |  |
| GENDEP | STAR*D | Improvement | 0.001 | 1429 | 0.00056 | 0.361 |
|  |  | Remission | 0.001 | 827 | 0.00199 | 0.130 |
| STAR*D | GENDEP | Improvement | 0.05 | 39471 | 0.00214 | 0.313 |
|  |  | Remission | 0.5 | 214280 | 0.00335 | 0.255 |

Secondly, we investigated whether genetic liability to MDD or schizophrenia predicted improvement in depressive symptoms, using meta-analyses from the Psychiatric Genomics Consortium (PGC) as discovery samples. Seven pharmacogenetic studies (including GENDEP and STAR*D) were used as independent target samples (3746 participants). Metaanalysis across studies (whole sample or citalopram/escitalopram treated patients) showed no predictive ability of genetic liability for MDD or for schizophrenia (Figure 2), with the most significant result being for schizophrenia PRS at P_T_<0.0001 (p=0.077). Across all P_T_, PRS for MDD showed p-values > 0.1 for the prediction of symptom improvement and regression coefficients explained less than 3% of the variance in symptom improvement. Results by-study at all P_T_ values are given in Supplementary Tables 1-2.

**Figure 2:**
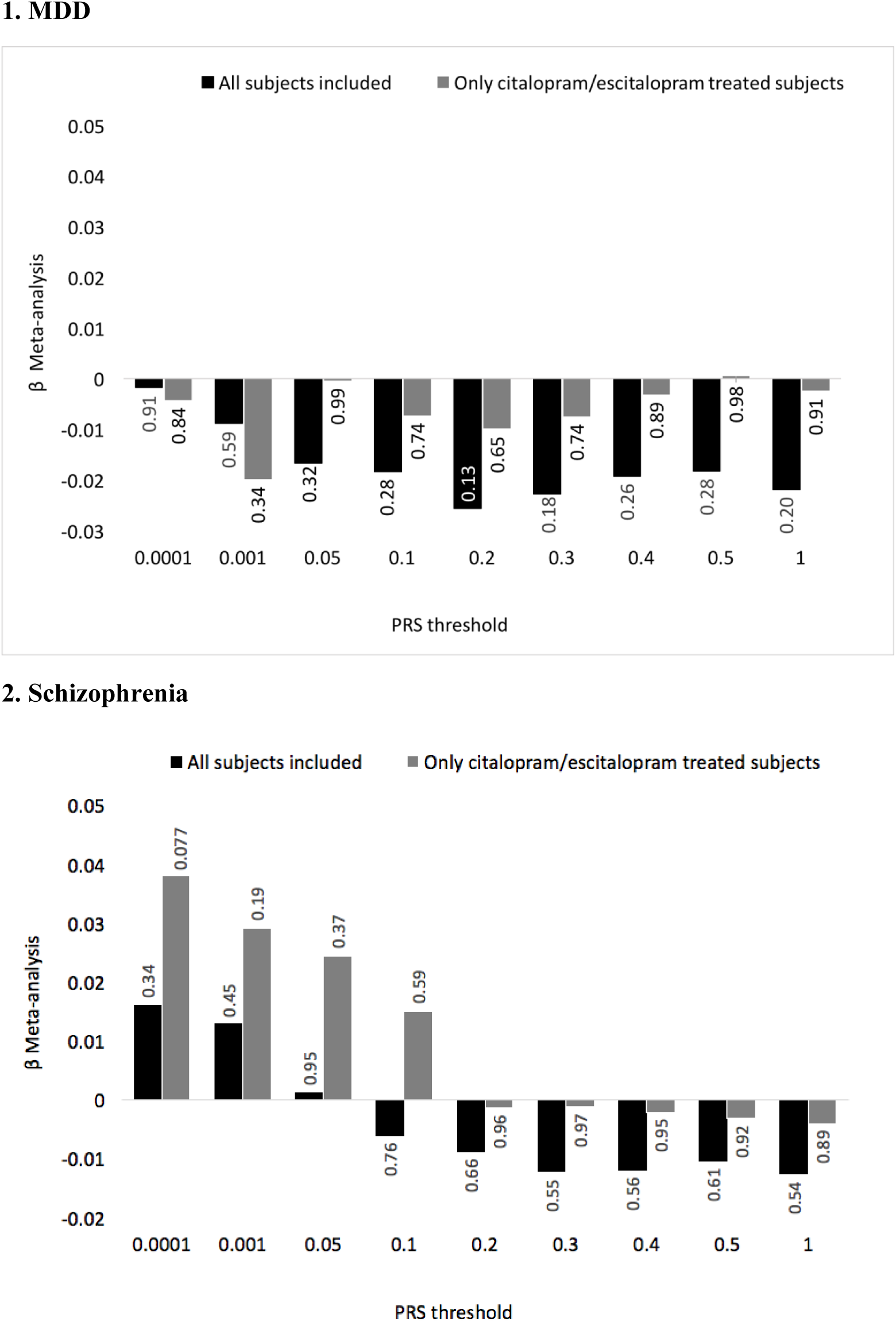
Meta-analysis of PRS effect sizes (β) in seven pharmacogenetic studies for (1) MDD and (2) schizophrenia PRS. Labels show p-values for meta-analyses at each p threshold.

## Discussion

In this study, we assessed whether the outcomes of antidepressant treatment may be predicted by PRS for (a) improvement and remission from an independent sample, (b) genetic liability to MDD, and (c) genetic liability to schizophrenia. Using each of the two largest available pharmacogenetic samples on antidepressant response (GENDEP and STAR*D) as baseline studies failed to predict antidepressant response in the other study. A previous study (GENDEP Investigators, MARS Investigators and STAR*D Investigators, 2013) found a small predicting ability of a PRS calculated in a meta-analysis of GENDEP-MARS studies in STAR*D, accounting for about 1.2% of the variance in outcomes in STAR*D. The present study was performed using individual datasets as discovery samples but increasing the number of genetic variants from ~ 1.2 million to ~ 7 million. PRS built from well-powered PGC studies for MDD and schizophrenia did not predict symptom improvement, either in individual pharmacogenetic studies, or in a meta-analysis. A previous analysis of PRS for bipolar disorder did not predict antidepressant response in STAR*D and the NEWMEDS studies, so this analysis was not repeated here (Tansey *et al.*, 2014).

This study represents the largest investigation of the PRS for antidepressant response to date, including the majority of currently available pharmacogenetic data on antidepressant response in MDD (3,746 participants from 7 studies). Both PGC discovery studies were well powered. The PGC schizophrenia study identified a genetic component accounting for approximately 7% of the liability to schizophrenia. MDD shares susceptibility genetic factors with schizophrenia (Cross-Disorder Group of the Psychiatric Genomics Consortium, 2013), suggesting a possible overlap between the genetics of schizophrenia and antidepressant response. The unpublished PGC MDD meta-analysis has a substantially increased sample size from the previous study (Ripke *et al.*, 2013b) as well as from the recent MDD GWAS from 23andme (Hyde *et al.*, 2016). MDD PRS comparable to the ones we calculated could not be constructed from the 23andme study since only SNPs with p<10^−5^ are publicly available.

Despite the extensive resources analysed, the power to detect predictions across study using PRS remained low for antidepressant response, although the power was adequate when we investigated common genetic liability with MDD and schizophrenia. The modest pharmacogenetic study sample sizes also precluded other whole-genome-approaches to estimate genetic correlation using GCTA or LD score regression (Yang *et al.*, 2011; Bulik-Sullivan *et al.*, 2015). A sample size ten-times larger would be required to achieve 80% for polygenic prediction between studies of antidepressant response. National registers and electronic medical records of large health care organisations could be used to achieve a study of this magnitude, but requires substantial resources for selection of appropriate subjects, phenotyping, DNA collection, genotyping and analysis. The power to detect common liability with psychiatric disorders was higher, but required the assumption of high genetic correlation.

Other limitations of the study arise from the differences in pharmacogenetic studies in characteristics of ascertainment, baseline severity, treatment, assessment of outcome and length of follow-up. We chose to focus on two largest studies (GENDEP, STAR*D) in test PRS for antidepressant response, to avoid adding multiple smaller studies where noise would outweigh signal. In the higher powered analysis assessing genetic component of MDD and schizophrenia, we included all available pharmacogenetic studies. Although there were substantial differences in the design of the studies, inclusion criteria were relatively similar and it was possible to establish comparable outcome measures. Ethnicity is also a possible cause of stratification in GWAS despite correction using ancestry-informative principal components.

We performed a single sub-analysis restricting to participants treated by citalopram or escitalopram, since escitalopram is the active isomer of citalopram (N=2308 participants) (Svensson and Mansfield, 2004). These analyses also failed to predict improvement of antidepressant symptoms or remission. Many further sub-hypotheses could be tested, for example, stratifying by sex, symptom dimensions, age, or severity. Recognising the need to balance a larger effect size in one subgroup against the smaller sample size and increased correction for multiple testing, we focussed on the key hypotheses (Traylor, Markus and Lewis, 2015).

The identification of individual genetic associations with antidepressant treatment response has been challenging, with no genome-wide studies identifying replicated signals for association (Uher *et al.*, 2010; Garriock *et al.*, 2010; Ising *et al.*, 2009). Since no major, single locus variants play a major role in treatment response, building polygenic predictors, which capture modest effects at multiple SNPs, may be a feasible alternative. STAR*D, GENDEP and other NEWMEDS studies show a strong polygenic component to the genetic architecture of response to antidepressants, with common genetic variation estimated to explain 42% of individual differences (SE = 0.180, p = 0.009) (Tansey *et al.*, 2013). With the decreasing costs of genotyping, and increasing access to such data, a PRS could form a powerful predictor response, and be of clinical value, as already seen in predicting disease risk (Chhibber *et al.*, 2014; Chatterjee, Shi and García-Closas, 2016). Other strategies, such as machine learning application to clinical and genetic variables in STAR*D and NEWMEDs studies showed some prediction based on both genetic and clinical characteristics, which was antidepressant specific (Iniesta *et al.*, 2016).

We selected here two reasonable polygenic hypotheses that (1) the genetic component of antidepressant response from a single study would transfer across studies, and (2) that genetic liability for psychiatric disorders would predict response to antidepressants. Neither of hypotheses could be confirmed in the currently available datasets and true polygenic component for antidepressant response would require much larger cohorts. Recent successes in uncovering the genetic component of psychiatric disorders are encouraging, but progress in uncovering the genetic component to treatment response remains slower. Expanded cohorts will be necessary to uncover the genetic architecture of antidepressant response, an essential step if precision medicine in depression is to become attainable.

## Acknowledgements

We thank the NIMH for access to data on the STAR*D study. We also thank the authors of previous publications in this dataset, and foremost, we thank the patients and their families who accepted to be enrolled in the study. Data and biomaterials were obtained from the limited access datasets distributed from the NIH-supported ‘‘Sequenced Treatment Alternatives to Relieve Depression’’ (STAR*D). The study was supported by NIMH Contract No. N01MH90003 to the University of Texas Southwestern Medical Center. The ClinicalTrials.gov identifier is NCT00021528.

We thank the University of Muenster, Germany, for having given the possibility of analysing their data on the MDD cohort that they collected between 2004 and 2006.

We also thank the authors of previous publications in these datasets, and foremost, we thank the patients who enrolled in these studies.

